# Bacteriophage SP01 Gene Product 56 (gp56) Inhibits *Bacillus subtilis* Cell Division by Interacting with DivIC/FtsL to Prevent Pbp2B/FtsW Recruitment

**DOI:** 10.1101/2020.02.06.938217

**Authors:** Amit Bhambhani, Isabella Iadicicco, Jules Lee, Syed Ahmed, Max Belfatto, David Held, Alexia Marconi, Aaron Parks, Charles R. Stewart, William Margolin, Petra Anne Levin, Daniel P. Haeusser

## Abstract

Previous work identified gp56, encoded by the lytic bacteriophage SP01, as responsible for inhibition of *Bacillus subtilis* cell division during its infection. Assembly of the essential tubulin-like protein FtsZ into a ring-shaped structure at the nascent site of cytokinesis determines the timing and position of division in most bacteria. This FtsZ ring serves as a scaffold for recruitment of other proteins into a mature division-competent structure permitting membrane constriction and septal cell wall synthesis. Here we show that expression of the predicted 9.3-kDa gene product 56 (gp56) of SP01 inhibits latter stages of *B. subtilis* cell division without altering FtsZ ring assembly. GFP-tagged gp56 localizes to the membrane at the site of division. While its localization permits recruitment of early division proteins, gp56 interferes with the recruitment of late division proteins, including Pbp2b and FtsW. Imaging of cells with specific division components deleted or depleted and two-hybrid analysis suggest that gp56 localization and activity depends on its interaction with mid-recruited proteins DivIC and/or FtsL. Together these data support a model where gp56 interacts with a central part of the division machinery to disrupt late recruitment of the division proteins involved in septal cell wall synthesis.

**IMPORTANCE:** Research over the past decades has uncovered bacteriophage-encoded factors that interfere with host cell shape or cytokinesis during viral infection. Phage factors that cause cell filamentation that have been investigated to date all act by targeting FtsZ, the conserved prokaryotic tubulin homolog that composes the cytokinetic ring in most bacteria and some groups of archaea. However, the mechanism of several identified phage factors that inhibit cytokinesis remain unexplored, including gp56 of bacteriophage SP01 of *Bacillus subtilis*. Here, we show that unlike related published examples of phage inhibition of cyotkinesis, gp56 blocks *B. subtilis* cell division without targeting FtsZ. Rather, it utilizes the assembled FtsZ cytokinetic ring to localize to the division machinery and block recruitment of proteins needed for the septal cell wall synthesis.

## INTRODUCTION

Most bacteria initiate cytokinesis through regulated assembly of the conserved tubulin-like GTPase FtsZ at the future site of division. FtsZ assembles into a toroidal array of treadmilling polymers that serve as a platform for recruitment of the cell division machinery, including enzymes needed for septal cell wall synthesis (1). Proper placement of the FtsZ ring in time and space is required to ensure that newborn cells reach adequate size and contain a full genetic complement. To achieve this, FtsZ assembly at mid-cell and subsequent division are highly precise, with less than a 1% margin of error, suggesting a highly regulated process (2, 3). Blocking FtsZ assembly prevents membrane invagination and septal cell wall synthesis, leading to filamentous, multinucleated cells and eventual cell death (4).

As a conserved protein that is essential for division in most bacteria, FtsZ is an appealing target of study for both physiologically relevant modes of its regulation and for potential development of novel antibiotics (5–7). Included among cellular regulators of FtsZ assembly are proteins encoded in regions of the *E. coli* genome that originally derived from phage, now turned inactive. Cells have co-opted several of these so-called cryptic phage genes for increased host fitness under particular conditions. These include *dicB* and *dicF* of cryptic phage Qin (aka phage Kim) and the *kilR* (*orfE*) gene of cyptic phage Rac (8). The RNA product of *dicF* binds to *ftsZ* mRNA to inhibit its translation (9), while the DicB peptide interacts with FtsZ inhibitor MinC (10) to target ring assembly independently of its normal regulator MinD, but dependent on ZipA (11). Transient division inhibition by cryptic DicB benefits the host by inhibiting phage receptor proteins ManYZ, enhancing immunity to bacteriophage lambda infection by up to 100-fold (12). The KilR peptide of Rac inhibits *E. coli* division through an unknown Min-independent mechanism that also causes increased loss of rod shape (13).

Functional bacteriophages also appear to encode factors that transiently block host cell division during infection. Expression of the 0.4 gene of T7 phage or *kil* of lambda phage both lead to *E. coli* cell filamentation through direct interference with FtsZ assembly by their protein products (14–16). In both cases, temporary inhibition of host cytokinesis by the phage prior to host lysis results in a subtle competitive advantage for the virus, although the specific nature of these advantages remains unclear.

Although all of the above factors come from phage that infect *E. coli*, it is likely that cytokinesis serves as a target for phage in the majority of other bacterial species as well. One reported example exists for the model Gram-positive species *Bacillus subtilis* and its lytic bacteriophage SP01, gene *56*, which lies in an operon comprising genes 58 through 56 (17). Expression of gene *56* alone, or in the context of the entire operon, leads to *B. subtilis* filamentation and death (18), but the mechanism behind this inhibition of cytokinesis is unknown.

The players involved in *B. subtilis* cytokinesis share commonalities with those involved in the *E. coli* division machinery, but several distinctions also exist (19, 20). Assembly of FtsZ at the membrane in *B. subtilis* involves interaction with the essential, well-conserved protein, FtsA, and with the non-essential protein, SepF, each of which contain a membrane-targeting sequence (21). The non-essential trans-membrane protein EzrA, an inhibitor of FtsZ assembly at the cell poles (22), also plays a separate role as part of the early division machinery by linking FtsZ indirectly to the membrane (23). Subsequent steps include recruitment of a trio of closely-interacting, membrane-spanning proteins: DivIB, FtsL, and DivIC. While *divIB* is dispensable in *B. subtilis* under laboratory conditions, both *ftsL* and *divIC* are essential for cytokinesis (24–26). Cellular levels of DivIB, FtsL, and DivIC are interdependent, closely linked through targeted proteolysis (27, 28). DivIB, FtsL, and DivIC cooperatively function as a complex to recruit the transpeptidase Pbp2B and the transglycosylase FtsW, both essential for septal cell wall synthesis (29).

Here, we characterize the activity of SP01 gene product 56 (gp56) in inhibition of *B. subtilis* cell division. We find that unlike all previously identified phage-derived inhibitors of cytokinesis, gp56 inhibits division independent of FtsZ. Instead, gp56 localizes to the *B. subtilis* division machinery in an FtsZ-dependent manner where it inhibits recruitment of later division components needed for septal cell wall synthesis. Our results suggest that localization of gp56 to the site of division involves interactions with essential division components FtsL and/or DivIC, and that disruption of their activity leads to reduced recruitment of Pbp2B and FtsW, resulting in cell filamentation and death.

## RESULTS

### Single copy, chromosomal expression of SP01 bacteriophage *GENE 56* inhibits *B. subtilis* cell division and sporulation

The previous report of division inhibition by gp56 utilized a multicopy plasmid, pPW19. To determine if gp56 is sufficient to inhibit division in single copy, we generated a strain, DPH102, in which gene *56* was expressed from a chromosomally encoded, IPTG-induced promoter at the amylase locus of *B. subtilis* strain JH642. Induction of gene *56* in DPH102 resulted in cell filamentation (Fig. 1A) similar to that previously reported in the *B. subtilis* CB-10 background using pPW18 (18). Consistent with division inhibition, staining the cell membrane with FM4-64 indicated that filaments were generally aseptate, with the exception of occasional septa at midcell. Such septa often appeared broader and ill-formed, consistent with a division block. Uninduced DPH102 cells were also slightly elongated (Fig. 1A), consistent with leaky expression of *gp56*.

**Figure 1.**
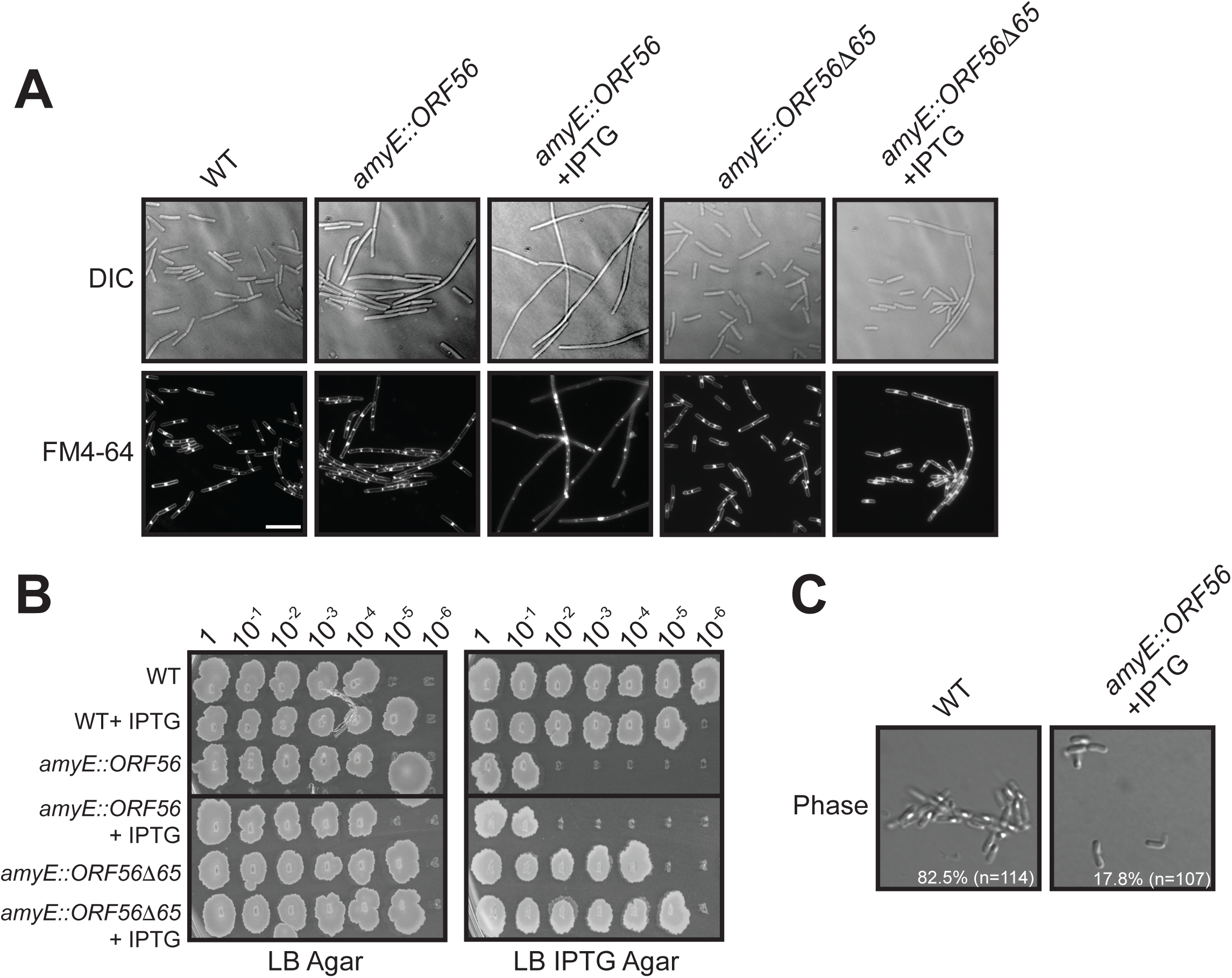
gp56 inhibits *B. subtilis* cell division. (**A**) Representative micrographs of live JH642 (WT), DPH102 (*amyE*::*56*), and DPH175 (*amyE*::*56Δ65*) cells from mid-log cultures (see Materials & Methods) grown in LB with or without 1 mM IPTG, as indicated, for induction of chromosomally-placed gene *56* or gene *56Δ65* at the *amyE* locus. Differential interference contrast (DIC) shows cells in brightfield (top row) and FM4-64 fluorescent staining (bottom row) shows cell membrane to differentiate between undivided, individual cell filaments and multiple, chained cells that form in the JH642 background with septal FM4-64 staining. Scale bar = 5 μm. (**B**) Spot titers of strains as in (A) taken from mid-log cultures grown in LB with or without 1mM IPTG, as indicated, and plated to either LB agar (left) or LB + 1mM IPTG agar (right). Dilution factor of spot titers from original cultures indicated on top. (**C**) Representative phase micrographs of JH642 (WT) and DPH102 (*amyE*::*56*) cells grown in Difco Sporulation Media, with or without 1mM IPTG as indicated. Spores appear as phase-bright dots. Scale as in (A). Percent quantification of total cell counts (n) showing a phase-bright spore for each strain is indicated inset.

For verification purposes, we additionally repeated published results from plasmid-borne gene *56* expression (18), but in our JH642 background. We transformed JH642 with pAP1 (pPW19 with the entire gene *58*-*56* operon) or pAP6 (pPW19 with gene *56* alone) to generate DPH176 and DPH3, respectively. As previously reported for the CB-10 background, IPTG induction of the operon, or gene *56* alone, both inhibited JH642 *B. subtilis* cell division indistinguishably, without altering cell growth or DNA replication/segregation (data not shown).

Serial dilution assays verified the lethality of long-term division inhibition, while also showing the reversibility of short-term division inhibition upon return to growth conditions without continued gene *56* induction (Fig. 1B). Spot plating of DPH102 mid-log culture dilutions to LB IPTG agar showed a four- to five-log reduction in long-term growth, both pre-grown in LB or LB IPTG liquid media. However, cells expressing gp56 survived comparably to the wild type parental strain if returned to media without IPTG.

*B. subtilis* typically grows for ∼4 to 5 mass-doublings in the absence of division (*e.g.* in the presence of gp56) before growth plateaus and cells entire a viable but non-culturable state (30). Consistent with the previous gp56 report (18), division inhibition by gp56 did not significantly alter these short-term *B. subtilis* growth rates. Under our growth conditions JH642 had a generation time (T_D_) of 29.0 +/- 1.1 minutes. Uninduced DPH102 had a similar T_D_ of 28.3 +/- 0.5 minutes, while induced DPH102 that express gp56 had a T_D_ of 30.7 +/- 2.2 minutes. Likewise, DNA staining verified that DNA replication and segregation appeared unaffected in DPH102 +/- IPTG (data not shown), consistent with previous reports of gp56 expression.

*B. subtilis* is well known for forming highly resistant endospores. A key step in sporulation is formation of an asymmetrically positioned septum that establishes two compartments: the larger mother cell and the small “forespore” (31). To determine if gp56 similarly impacts asymmetric division, we assessed its impact on spore formation. Phase-contrast microscopy of JH642 cells plated to DSM agar showed ∼83 – 90% of cells with fully formed phase-bright spores (Fig. 1C). Consistent with leaky production of gp56 being sufficient to inhibit asymmetric division, only ∼40% in % uninduced DPH102 cells. Only ∼18% of DPH102 cells showed signs of a phase-bright spore upon gp56 expression (Fig. 1C). Moreover, of that minority, most appeared to be bands of phase brightness rather than the typical round shape of a properly formed spore.

### The carboxyl-terminal domain of gp56 is essential for division inhibition

To identify regions of gp56 required for division inhibition, we serially passaged DPH3 (JH642 pAP6) on LB IPTG agar to isolate suppressors that grow in the presence of gp56. Plasmid sequencing revealed the overwhelming majority of such isolates had either promoter mutations or nonsense mutations early in the gene 56 coding sequence. One isolate (gene *56Δ65*), however, contained a nonsense mutation at codon 65 of the gene, truncating the predicted gp56*Δ*65 product by 15 of 79 residues. Analysis of the primary sequence of gp56 by SMART (32) predicts that residues 37 – 59 form a transmembrane domain, with a highly favored orientation prediction by TMpred (33)where its amino terminus faces the cytoplasm and its carboxyl-terminus faces extracellularly.

To assess the phenotype of gp56*Δ*65 comparably to the WT, we sub-cloned the gene *56Δ65* allele from its isolated plasmid (pDH89) and placed it under IPTG-inducible control at the *amyE* locus of JH642 to generate DPH175. As expected, single-copy, chromosomal expression of gp56*Δ*65 failed to cause cell filamentation (Fig. 1A) or lethality (Fig. 1B). This suggests that gp56*Δ*65 has lost its ability to inhibit *B. subtilis* cell division, and implicates the predicted extracellular C-terminus of gp56 as essential for mediating that inhibitory phenotype.

### Division inhibition by gp56 is independent of FtsZ ring formation

Previously characterized bacteriophage division inhibitors function by reducing cellular FtsZ concentrations (*e.g. dicF* (9)) or directly antagonizing its assembly (*e.g.* l *kil* (16, 34)). To determine if the same is the case for gp56, we used immunofluorescence microscopy (IFM) to localize FtsZ in wild type, DPH102, and DPH175 cells in the presence an absence of inducer.

As expected for the JH642 wild type background, the majority of cells displayed a single FtsZ ring at mid-cell (Fig. 2A), with an average cell length of 4.1 μm and width of 1.3 μm (Figs. 2B & 2C). Similar to wild-type cells, uninduced DPH102 cells likewise contained single, medial FtsZ rings (Fig. 2A) with an average cell length of 4.5 μm and width of 1.4 μm (Figs 2B & 2C). As expected, expression of gp56 in DPH102 resulted in a division block, increasing average cell length to ∼18.6 μm (Fig. 2B). The average length measurement of these filaments is likely an underestimate, as many filaments extended past the micrograph field of view. Somewhat surprisingly, we observed regularly spaced FtsZ rings along the length of DPH102 cells cultured in the presence of IPTG (Fig. 2A). As expected, almost all DPH175 cells contained a single FtsZ ring at midcell (Fig 2A) regardless of gp56*Δ*65 expression. Additionally, DPH175 fixed cell lengths were not altered by expression of the truncated gp56*Δ*65 (4.2 and 3.9 μm, respectively) (Fig 2B). Uninduced DPH175 cells did have a slightly larger average cell width (1.5 μm) compared to induced conditions (1.3 μm) or that of the other strains. While a statistically significant increase by analysis of variance, it is likely an artifact of the fixation process, as similar changes in width are not observed in live cells.

**Figure 2.**
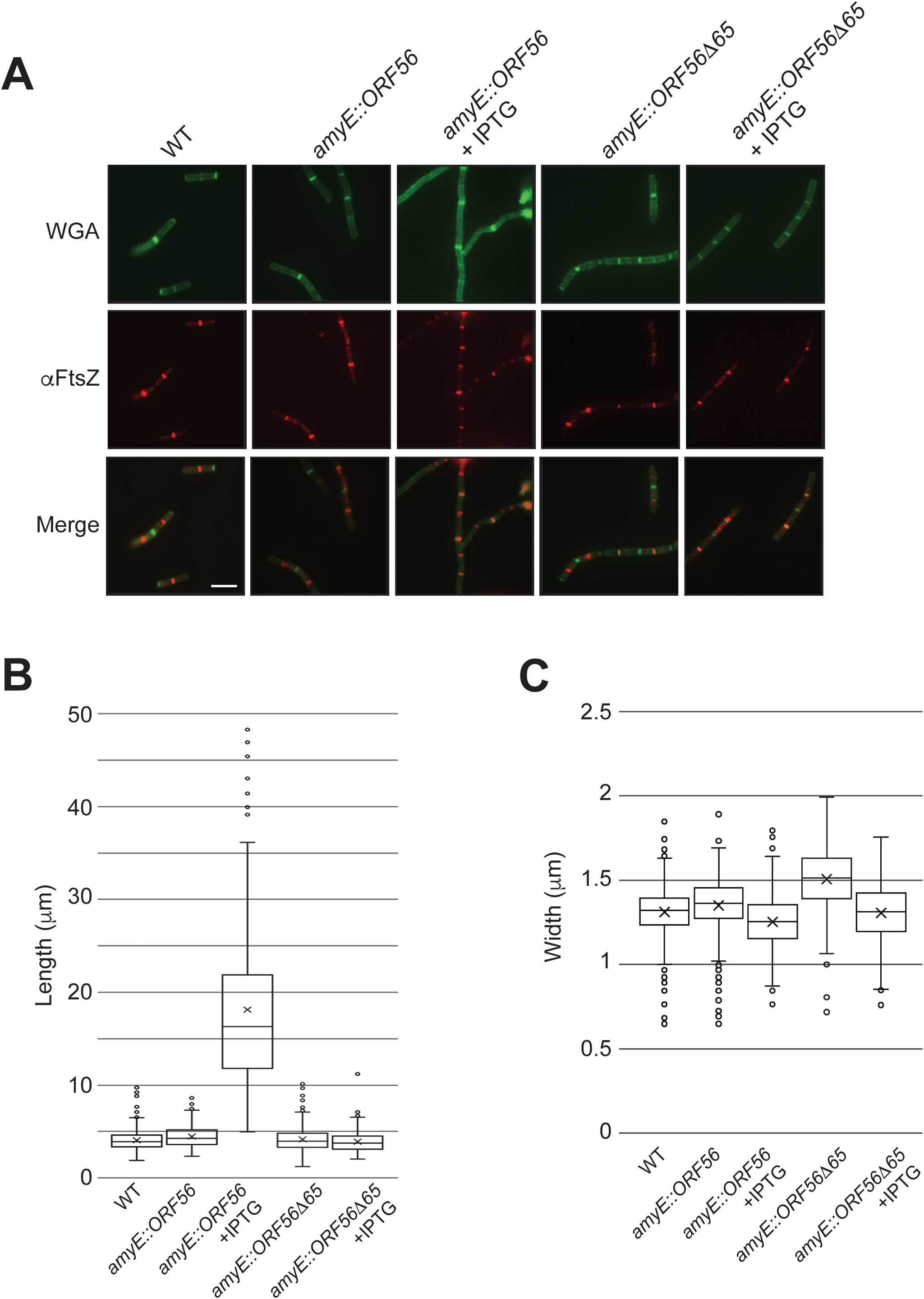
gp56 does not inhibit FtsZ ring assembly in *B. subtilis*. (**A**) Representative false-colored immunofluorescence micrographs of glutaraldehyde/paraformaldehyde-fixed JH642 (WT), DPH102 (*amyE*::*56*), and DPH175 (*amyE*::*56Δ65*) cells taken from mid-log cultures with or without 1mM IPTG as indicated. Fluorescent Wheat germ agglutinin (WGA) staining (top row) shows cell wall, *α*FtsZ (middle row) shows signal from fluorescent-conjugated secondary antibody to primary antibody against FtsZ. Bottom row shows a merge of the WGA and *α*FtsZ signals. (**B**) Scatter box plots of cell length (μm) quantification (n > 100) of strains from (A). Box borders denote upper and lower quartiles, with horizontal line in box depicting the median and X depicting the mean. Whiskers show upper and lower deviation of data. (**C**) Scatter box plots of cell width (μm) quantification (n > 100) of strains from (A), with data shown as in (B).

Although DPH102 filaments formed by gp56 expression still contain FtsZ rings, it is possible they assemble less efficiently. Calculations of length/ FtsZ ring (LR) ratios in DPH102 filaments compared to cells of normal length allow determination of any alteration in FtsZ ring frequency/stability in the presence of gp56. Un-filamented cells had an average L/R ratio of 4.2. In contrast, filamentous DPH102 cells had an average L/R ratio of 7.2. This suggests that while gp56 does not abolish FtsZ ring assembly, it does somewhat reduce them, likely through prolonged filamentation leading to disassembly of preformed FtsZ rings rather than any reduction in their *de novo* assembly. Regardless, such a modest increase in L/R ratio would not explain the observed extreme filamentation.

### *gp56* interferes with recruitment of essential late division proteins to the site of division

Together, the preceding data indicate that gp56 acts to block cell division at a step following FtsZ ring formation. Preventing recruitment of other essential proteins of the division machinery to FtsZ, such as enzymes involved in septal peptidoglycan synthesis, also produces a filamentous cell phenotype. To identify the target of gp56, we assayed recruitment of individual GFP-tagged division components in the JH642 *amyE*::*P_spac_-56* (DPH102) background, in the absence and presence of IPTG inducer.

Our resulting data indicate that gp56 mediated division inhibition occurs through disruption of late protein localization. Green fluorescent protein (GFP) fusions to relatively early localized division proteins EzrA (DPH111) and DivIB (DPH97) exhibited regular staining patterns in filaments formed from gp56 expression, comparable to cells lacking gp56 (Fig. 3). In contrast, DivIVA-GFP (DPH371), one of the last proteins recruited during active septation as new daughter cell poles form, exhibited diffuse cytoplasmic staining in filaments formed from gp56 expression.

**Figure 3.**
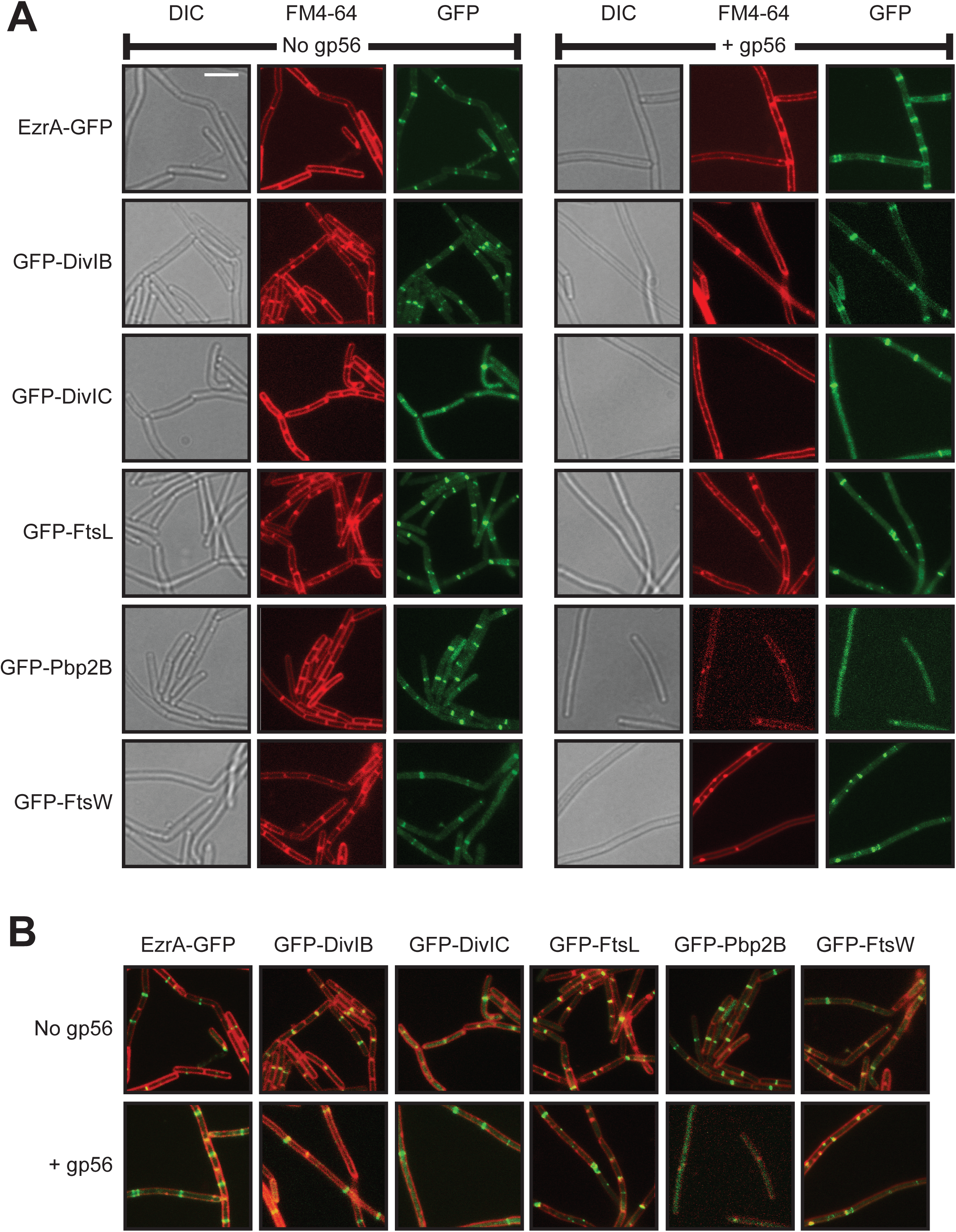
gp56 prevents recruitment of late, essential division proteins needed for *B. subtilis* septal cell wall synthesis. (**A**) Representative false-colored fluorescent micrographs of live cells of strains expressing EzrA-GFP (PL847 and DPH111), GFP-DivIB (DPH79 and DPH97), GFP-DivIC (DPH617 and DPH618), GFP-FtsL (DPH1108 and DPH584), GFP-Pbp2B (DPH414 and DPH415), or GFP-FtsW (DPH408 and DPH409) in the absence (left) or presence (right) of gp56. Each field of view includes a DIC image showing cells in brightfield, FM4-64 showing fluorescently stained membranes, and GFP fluorescent signal showing localization of the indicated fusion construct. Scale bar = 5 μm. (**B**) Merged panels, in same scale, of WGA and GFP signal of from (A).

We next constructed *gfp*-tagged fusions to *ftsW* and *pbp2b* and cloned these at the amylase locus under xylose control (DPH387 and DPH400, respectively). These strains were then transformed with pPW19 or pAP6 to assess potential gp56 effects on GFP-FtsW (DPH408 and DPH409) or GFP-Pbp2B (DPH414 and DPH415) localization. As later recruited proteins, fewer cells in the population normally contain localized GFP-FtsW or GFP-Pbp2B at any given time, compared to early recruited proteins that remain localized for most of the cell cycle. Nonetheless, in the absence of gp56, visible GFP-FtsW and GFP-Pbp2B bands were apparent at mid-cell in multiple cells within a given field of view. In contrast, in the presence of gp56 no cytoplasmic bands of GFP-Pbp2B were observed, and the only GFP-FtsW bands visible were less uniform in shape (Fig. 3) and frequently overlapped with blebs of improperly formed septal cell wall visible through fluorescent WGA staining (Fig. 3B)

The diminished recruitment of GFP-FtsW and GFP-Pbp2B by gp56 suggested that the phage peptide might interfere with formation of the DivIB-FtsL-DivIC complex, which cooperatively recruit FtsW and Pbp2B for cell wall septum formation (29). To test this possibility, we constructed *gfp*-tagged fusions to *ftsL* and *divIC* and inserted them at the amylase locus under xylose control (DPH579 and DPH614, respectively). These strains were then transformed with pPW19 or pAP6 to assess potential gp56 effects on GFP-FtsL (DPH1108 and DPH584) or GFP-DivIC (DPH617 and DPH618) localization.

Upon their induction, both GFP-FtsL and GFP-DivIC each demonstrated localization to mid-cell in the absence of gp56, as expected. Additionally, GFP-FtsL and GFP-DivIC still were capable of localizing as cytoplasmic bands within cell filaments formed from gp56 expression. However, this localization of FtsL and DivIC appeared reduced in the presence of gp56, compared to its absence, and FtsL in particular showed a striking occasional pattern of disrupted localization in helices or blobs (Fig. 3).

### gp56 co-localizes with the division machinery in a manner that requires FtsZ, FtsA, as well as DivIC and/or FtsL

An inhibitor of bacterial cell division that acts after FtsZ ring assembly might localize to midcell via interactions with components of the division machinery to prevent recruitment of relatively later components. To determine if this is true for gp56 and the observed loss of FtsW and Pbp2B recruitment in its presence, we constructed a chromosomal fusion of *gfp* to gene *56* at the *amyE* locus under control of xylose (DPH50). We chose an N-terminal GFP tag based on the prediction that the C-terminal end of the gp56 peptide would be extracellular and the N-terminal end intracellular. As a control we also constructed a strain with a similar *gfp* fusion to the inactive gene *56Δ65* allele (DPH170).

Induction of *gfp-56* resulted in cells with a clear band of GFP fluorescence at mid-cell, consistent with GFP-gp56 localization to the divisome. In contrast, induction of the inactive truncated gp56 control showed poor to little localization (Fig. 4A). Notably, the addition of the GFP tag to the gp56 peptide does seem to interfere with the peptide’s activity in blocking cell division, as cells expressing *gfp-56* did not significantly filament.

**Figure 4.**
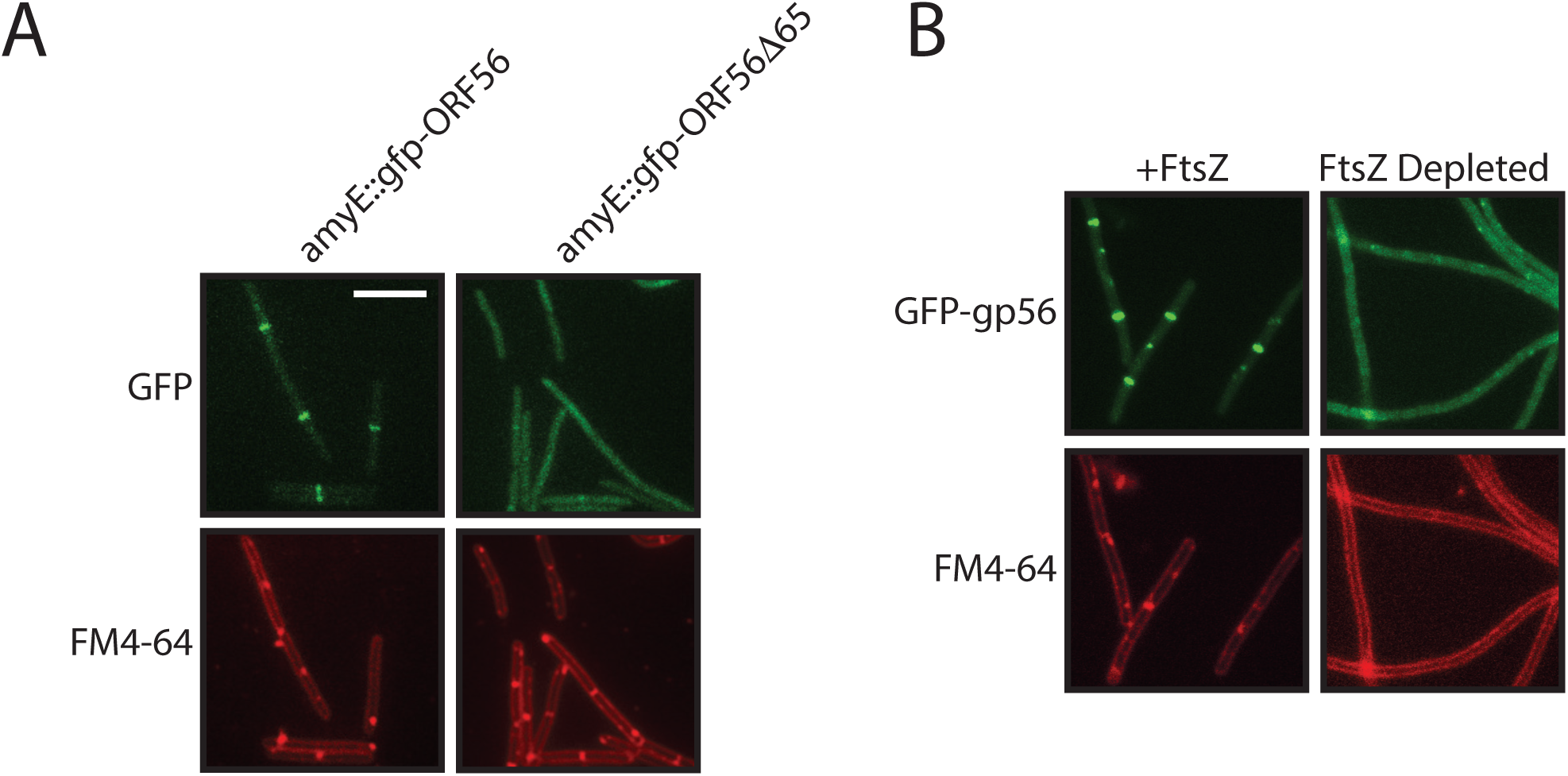
GFP-gp56 localizes to the *B. subtilis* site of division in an FtsZ-dependent manner. (**A**) Representative false-colored fluorescent micrographs of live DPH50 (*amyE*::*gfp-56*), and DPH170 (*amyE*::*gfp-56Δ65*) cells taken from mid-log cultures with 0.1% xylose present to induce fusion protein expression. GFP signal (top) shows fusion protein localization and FM4-64 fluorescent signal (bottom) shows cell membranes. Scale bar = 5 μm. (**B**) Representative false-colored fluorescent micrographs of live DPH504 (*ftsZ*::*spc xylA*::*tet thrC*::*P_xyl_*-*ftsZ amyE*::*P_spac_*-*gfp-56*) cells taken from mid-log cultures with 1 mM IPTG present to induce GFP-gp56 in the presence (left) or absence (right) of xylose to express/deplete FtsZ. GFP signal (top) shows GFP-gp56 localization and FM4-64 fluorescent signal (bottom) shows cell membranes. Scale as in (A).

To determine if gp56 localization was dependent on FtsZ or downstream components of the divisome we transformed the *amyE*::*P_xyl_*-*gfp*-*56* region of the chromosome into strains that contain deletions, or permit depletion, of specific components of the cell division machinery. Depletion of *ftsZ* (DPH504) abolished GFP-gp56 localization to midcell (Fig. 4B), confirming that observed GFP-gp56 bands reflect localization to the site of division in a manner requiring the foundational assembled FtsZ.

We next sought to address what other components – if any in addition to FtsZ – were involved in recruitment of gp56 to its site of activity at the divisome. Not surprisingly, depletion of *ftsA* (DPH503) also led to the loss of almost all GFP-gp56 localization. However, deletion of either *divIB* or *ezrA* (DPH55 and DPH177, respectively) did not result in any alteration of GFP-gp56 localization compared to that seen in the wild-type background (Fig. 5).

**Figure 5.**
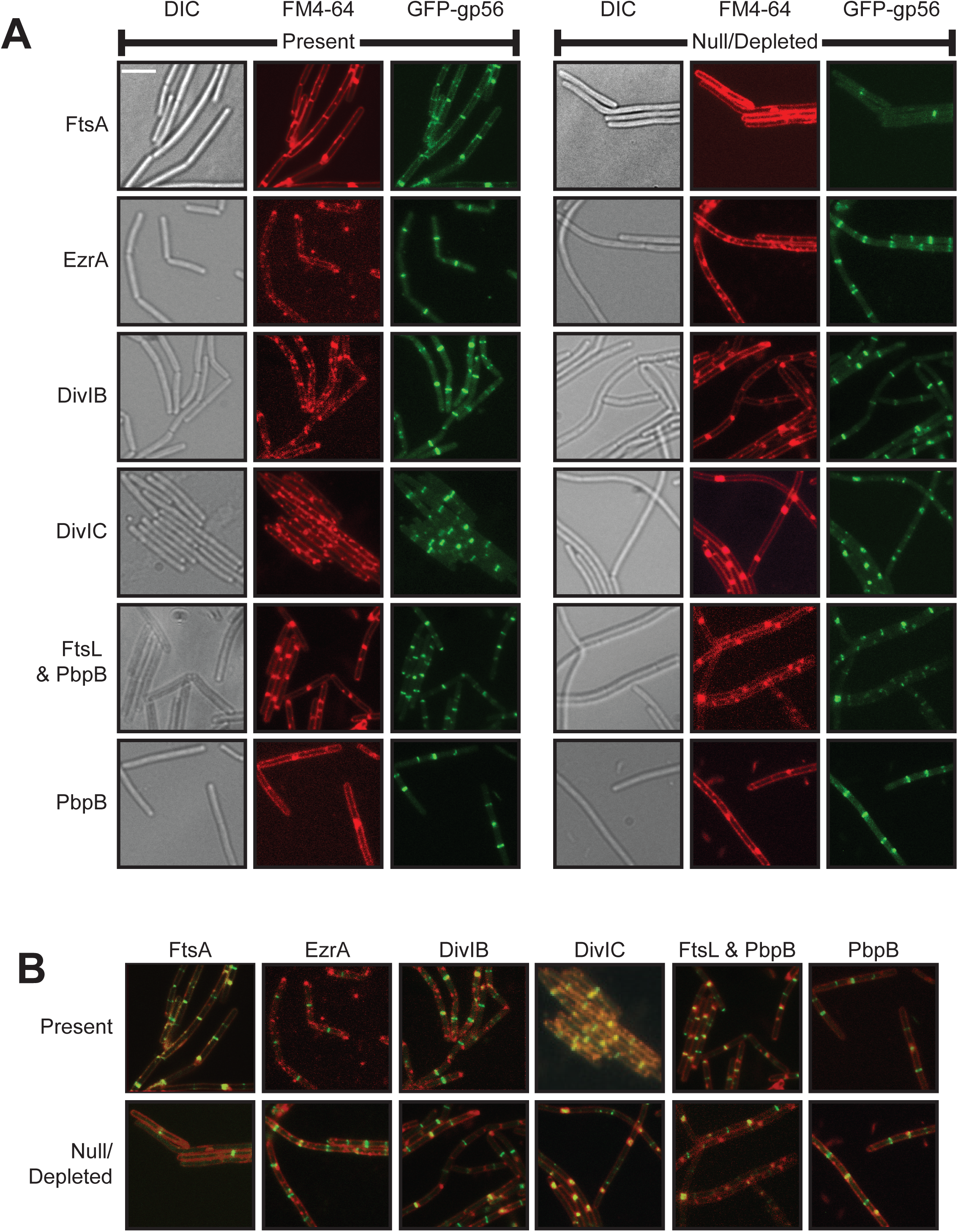
GFP-gp56 localization to the *B. subtilis* site of division requires DivIC/FtsL. (**A**) Representative false-colored fluorescent micrographs of live cells taken from mid-log cultures containing (left) or missing (right) FtsA (DPH503), EzrA (JH642 or DPH55), DivIB (JH642 or DPH177), DivIC (DPH302), FtsL and Pbp2B in combination (DPH1121), or Pbp2B alone (DPH1119) through deletion/depletion. Each field of view includes a DIC image showing cells in brightfield, FM4-64 showing fluorescently stained membranes, and GFP fluorescent signal showing localization of GFP-gp56. Scale bar = 5 μm. (**B**) Merged panels, in same scale, of WGA and GFP signal of from (A).

Given these results and the observed loss of Pbp2B and FtsW localization in the presence of gp56, this leaves DivIC and FtsL as potential key factors that may contribute to gp56 recruitment to the divisome. Depletion of *divIC* resulted in a loss of the majority of GFP-gp56 fluorescent bands (Fig. 5). Instead, GFP-gp56 signal mostly appeared at the periphery of cells (membrane localization) and to unproductive septal patches that appeared due to the filamentation of cells depleted for *divIC* (Fig. 5B). This argues that DivIC does play a significant role in gp56 recruitment to the divisome. Likewise, depletion of *ftsL* (DPH1119) led to a loss in GFP-gp56 localization (Fig. 5). Notably, however, this strain does not allow the depletion of *ftsL* alone, but only in conjunction with *pbpB* (encoding Pbp2B), within the context of the *yllB*-*ylxA*-*ftsL*-*pbpB* operon. Because gp56 appears to prevent Pbp2B localization to the divisome, we would predict that depletion of *pbpB* alone should not lead to any loss of GFP-gp56 recruitment to the divisome. Indeed, following depletion of *pbpB* alone (DPH1121), GFP-gp56 retained its normal localization (Fig. 5). This argues that the observed loss of its localization following depletion of *ftsL* and *pbpB* together likely stems from the lack of FtsL specifically.

### DivIC and FtsL interact with gp56 by bacterial two-hybrid assay

Observed disruptions in normal DivIC and FtsL localization in the presence of gp56, coupled with the observed dependence of gp56 localization on the presence of DivIC and FtsL suggests that these three components might directly interact at the divisome. To test for the potential interactions between these proteins, we cloned *divIC*, *ftsL*, gene *56*, and gene *56Δ65* each into pKT25 and gene *56* into pUT18C for analysis by the bacterial two-hybrid assay (35). We combined each pKT25 construct together with pUT18C-gene *56* in the DHM1 reporter strain and plated the resulting strains to appropriate media. The results of these bacterial two-hybrid analyses indicate that gp56 is capable of strong self-interaction as well as interaction with both FtsL and DivIC. In contrast, interaction with gp56*Δ*65 is absent (Fig. 6A).

**Figure 6.**
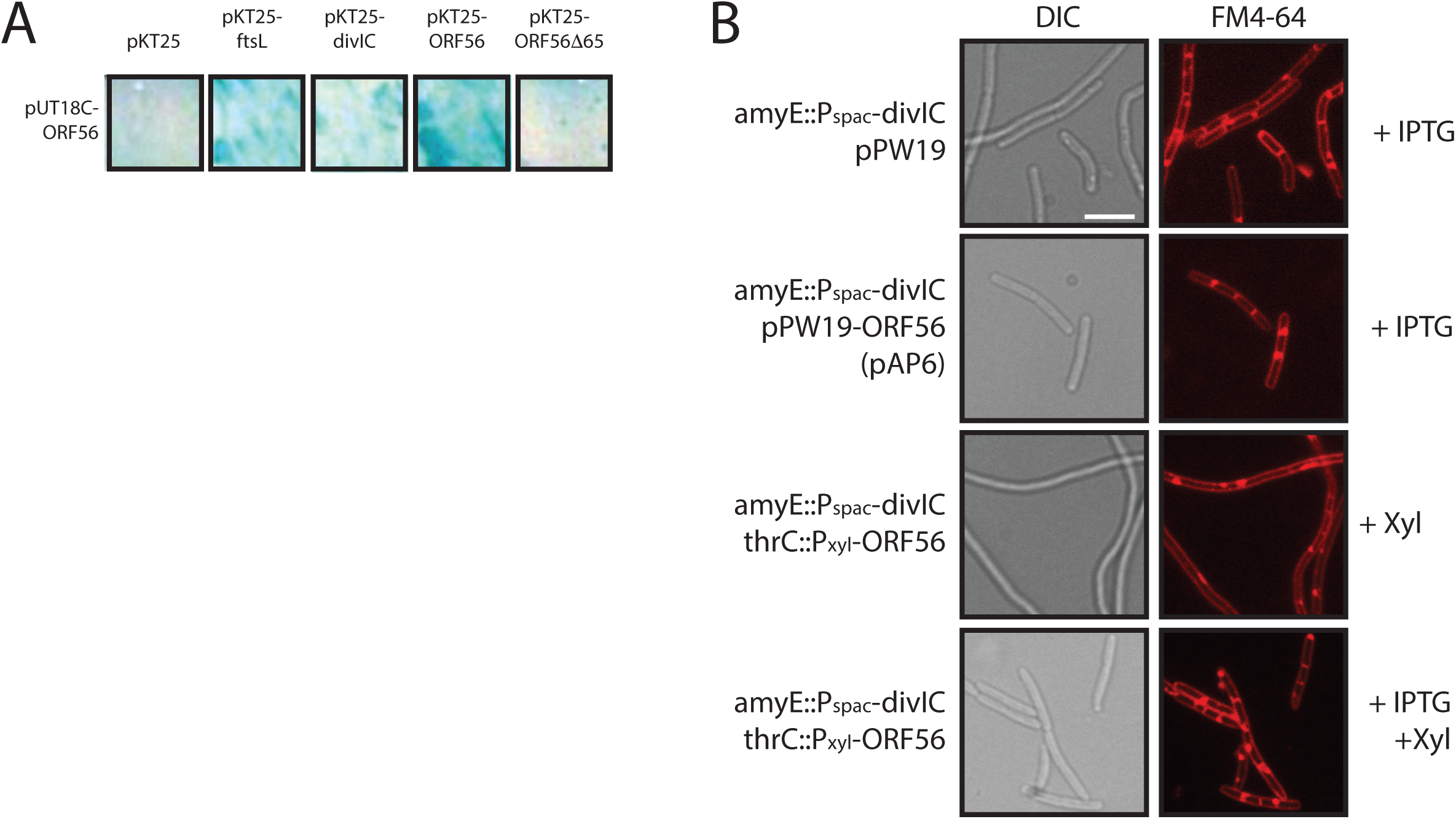
SP01 gp56 interacts with itself and *B. subtilis* DivIC or FtsL, and its activity is suppressed by *divIC* over-expression. (**A**) Representative patched growth of DHM1 background *E. coli* strains DPH183 (gp56 alone), DPH164 (gp56 and FtsL), DPH165 (gp56 and DivIC), DPH166 (two copies of gp56), and DPH167 (gp56 and gp56*Δ*65) bacterial two-hybrid strains containing pUT18C-*56* and indicated pKT25 constructs on LB Amp^100^ Kan^50^ agar plates with 1 mM IPTG and 50 g/ml X-Gal. (**B**) Representative false-colored fluorescent micrographs of live cells taken from mid-log cultures with DIC (left) showing cells in brightfield and FM4-64 (right) showing fluorescently stained membranes. DPH660 (*amyE*::*P_spac_-divIC pPW19*) and DPH661 (*amyE*::*P_spac_-divIC pPW19-56*) grown with 1 mM IPTG to overexpress *divIC* and express gene *56* from pPW19 (top two rows). DPH1152 (*amyE*::*P_spac_-divIC thrC*::*P_xyl_-56*) grown with 0.1% xylose to express gene *56* in the absence (third row) or presence (bottom row) of 1 mM IPTG to overexpress *divIC*. Scale bar = 5 μm.

### DivIC overexpression supports the model that gp56 interacts with it to inhibit cell division

The above results are consistent with a model where gp56 localizes to the divisome through interactions with DivIC and/or FtsL (in turn dependent on FtsZ and FtsA). The interaction of DivIC and FtsL with gp56 partially disrupts their normal localization and thereby presumably compromises their normal functions, leading to a loss of Pbp2B and FtsW recruitment and thereby cell filamentation. We hypothesized that if gp56 interactions with DivIC and/or FtsL are disrupting their normal functions, then perhaps overexpression of either of these components would dilute out the inhibitory effects of gp56 and restoring cell division.

While simultaneous overexpression of *ftsL* and *pbp2B* did not have any effect on gp56 inhibitory activity (data not shown), we found that overexpression of *divIC* through two separate constructs did suppress gp56 inhibition of cell division (Fig. 6B). In the first construct, we utilized a strain with an IPTG-inducible second copy of *divIC* at the amylase locus and transformed it with either pPW19 or pAP6. For the resulting strains (DPH660 and DPH661), IPTG addition simultaneously induces overexpression of *divIC* and gene *56* in pAP6. In contrast to the filamentation seen normally upon gp56 expression from pAP6, no cell filamentation occurred with the simultaneous overexpression of *divIC* (Fig. 6B), suggesting that extra DivIC is protective against gp56 activity. For the second construct, we the divIC overexpression background with chromosomal expression of gene *56* from the *thrC* locus through xylose induction (DPH1152). When xylose alone was added to the resulting strain, gp56 expression led to cell filamentation as expected. However, inclusion of IPTG in addition to the xylose allowed *divIC* overexpression and led to rescue from the gp56-mediated block in cell division (Fig. 6B). Together these data support the model in which gp56 interferes with DivIC to block further divisome component recruitment, but that additional cellular DivIC can dilute out these effects.

## DISCUSSION

The results reported here demonstrate that the inhibition of *B. subtilis* cell division by gp56 of bacteriophage SP01 is exerted downstream of FtsZ ring assembly. FtsZ rings assemble at regular intervals along undivided *B. subtilis* filaments in the presence of gp56, but these rings do not initiate visible constriction or septation. Expression of division components fused to GFP demonstrates that many still localize in the presence of gp56, but that Pbp2B and FtsW recruitment becomes dramatically reduced. Moreover, mid-cell bands of FtsL and DivIC are slightly reduced or disrupted in the presence of gp56. GFP-tagged gp56 demonstrates FtsZ- and FtsA-dependent localization to mid-cell, and its apparent recruitment to the division machinery also requires DivIC and FtsL. Consistent with these results, gp56 shows interaction with itself, FtsL, and DivIC by bacterial two-hybrid analysis, and overexpression of *divIC*, but not *ftsL* can suppress gp56-mediated division inhibition.

Together, our results suggest a model (Fig. 7) in which gp56 localizes to the division complex in an FtsAZ-dependent manner through interactions with DivIC/FtsL, where its presence interferes with normal DivIB/FtsL/DivIC activity and proper recruitment of Pbp2B and FtsW.

**Figure 7.**
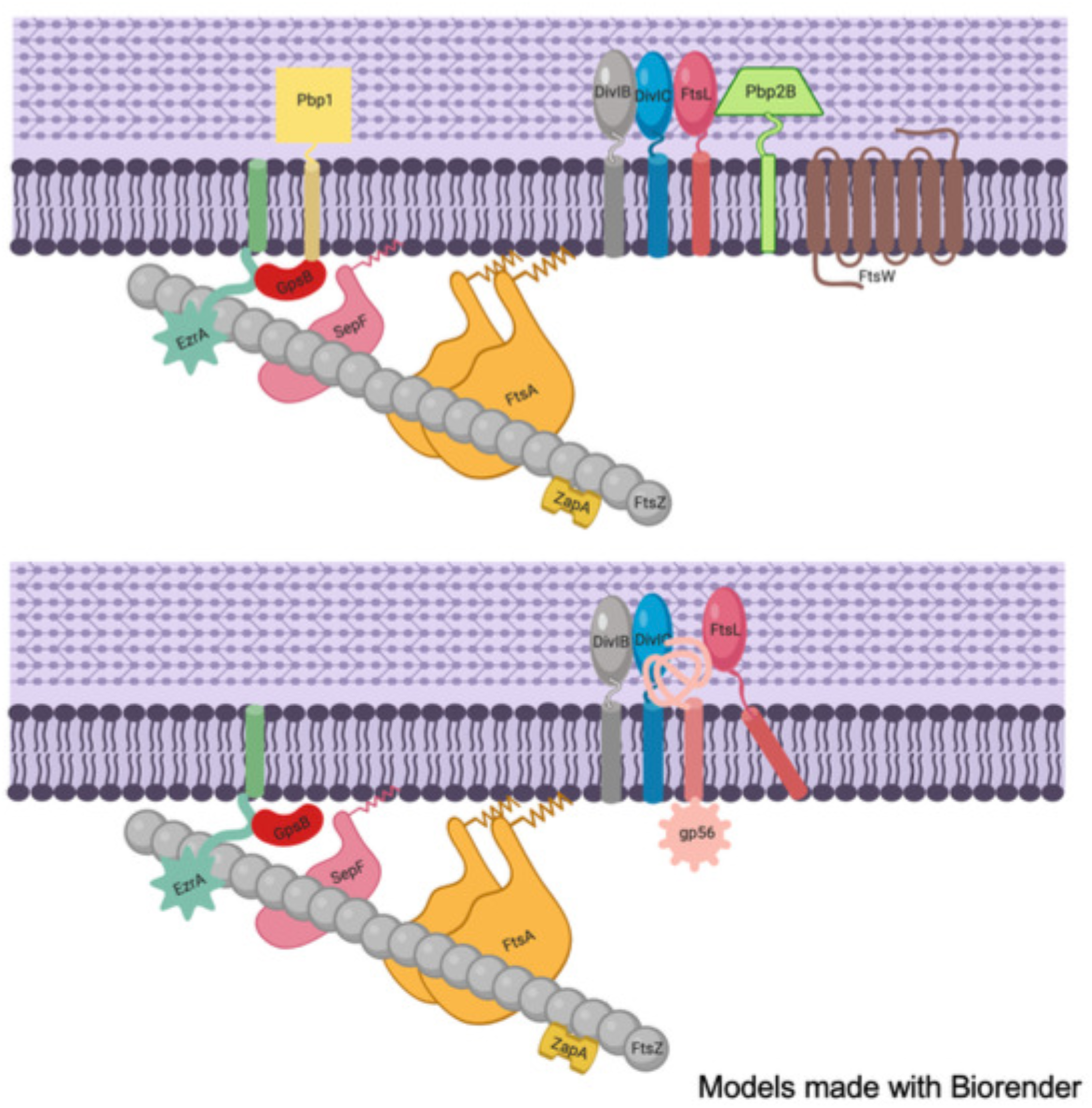
SP01 gp56 interacts with *B. subtilis* DivIC/FtsL to disrupt the division machinery and prevent recruitment of the Pbps and FtsW that are essential for septal cell wall synthesis. (**A**) In normal conditions the *B. subtilis* division machinery assembles into a cytokinesis-competent apparatus allowing for proper membrane constriction and septal wall synthesis. FtsZ polymers assemble at the membrane through interactions with membrane-associated FtsA, SepF, and EzrA. Stabilized by non-essential ZapA assembled FtsZ and its membrane-associated partners allow recruitment of DivIB/FtsL/DivIC complexes that in turn help recruit Pbp2B and FtsW, while non-essential GpsB and EzrA help recruit Pbp1. (**B**) In the presence of SP01 gp56, the phage peptide interacts with DivIC/FtsL, mildly disrupting their normal localization and thereby preventing the normal recruitment of Pbp2B, FtsW, Pbp1, and other late division proteins (*e.g.* DivIVA, not shown). While lack of this late recruitment would also normally lead to rapid loss of the DivIB/DivIC/FtsL complex, its interaction with gp56 prevents their loss, effectively freezing the division machinery at a mid-assembled state unable to constrict or build septal cell wall, thereby leading to cell filamentation and death. Cartoon models generated through Biorender.

SP01 gp56 joins an expanding cast of phage-derived factors that inhibit host cytokinesis, including Kil of bacteriophage lambda (16, 34), gp0.4 of bacteriophage T7 (14, 36), and elements of defective prophage like DicB (12) or *dicF* of Qin (9) and Kil of Rac (13). Notably, however, SP01 gp56 represents the first phage factor described to not target FtsZ directly, or indirectly, in its inhibition of cytokinesis. Instead, gp56 has likely evolved to target division machinery maturation by blocking recruitment of septal cell wall synthesis enzymes. A similar mechanism appears to have evolved in *B. subtilis* with the SOS-induced division inhibitor YneA, which likewise blocks cytokinesis downstream of FtsZ ring assembly (37).

In the case of SP01 gp56, this strategy also allows the bacteriophage to potentially make use of a nonfunctional division machinery foundation for localization of its own factors, at least gp56. A similar case occurs with p1 of *B. subtilis* bacteriophage *ϕ*29, which localizes to assembled FtsZ at mid-cell to promote phage particle assembly (38). Localization of p1 requires FtsZ, but not Pbp2B, however whether it interacts directly with FtsZ or utilizes another division protein for its localization like gp56 is unknown. Regardless of its precise interactions, *ϕ*29 p1 recruitment to the division machinery only modestly interferes with *B. subtilis* cytokinesis (38), unlike the total block in division caused by SP01 gp56.

Within the SP01 genome, gene *56* is found at the end of an operon with genes *58* and *57* (17, 18), two genes whose products also lack any homologs in databases, and whose function is unknown. It is possible that the products of these other genes colocalize along with gp56 to the FtsZ ring to carry out an activity for SP01 processing together, similar to that seen with *ϕ*29 p1, that transiently delays cytokinesis sufficiently to prevent division septum formation from interfering with assembling viral particles.

Analogous temporary division blocks also occur within the *B. subtilis* host, such as during the aforementioned YneA-mediated SOS response to permit DNA repair (37), or during the transition of from vegetative growth to sporulation via RefZ activity on FtsZ (39, 40). It also occurs during the DNA recombination events that accompany the developmental stress response of *B. subtilis* natural competence, where the peptide Maf is produced upon DNA uptake, and directly and transiently inhibits FtsZ assembly to permit uninterrupted genome maintenance (41).

The previous study (18) identifying SP01 gp56 as an inhibitor of *B. subtilis* cytokinesis demonstrated that a temporary block in division does occur during the SP01 infective process prior to host cell lysis. However, SP01 lacking gene *56* displays no apparent phenotypic defect in burst size or latency under laboratory conditions (18). It still remains possible, however, that gp56-mediated cytokinetic blocks give subtle competitive advantages to SP01 under particular growth conditions by preventing cells from dividing over phage particles in the process of assembly. Additionally, given that gp56 also interferes with maturation of the asymmetric FtsZ ring formed during sporulation, that developmental pathway may represent a situation where gp56 becomes more important for SP01 infective success.

Beyond the roles for gp56 in SP01 biology, its apparent interactions with FtsL/DivIC make it a potential tool for further study of the role that these proteins play in *B. subtilis* cell division. Normally, loss of Pbp2B or FtsW localization to the division machinery perturbs ‘back-recruitment’ of the ternary complex of DivIB/FtsL/DivIC, where they too become delocalized (27–29). However, in the presence of gp56, loss of Pbp2B and FtsW does not result in that perturbation, presumably because FtsL/DivIC interactions with gp56 help protect them from delocalization and/or proteolysis, even while rendering them at least partially dysfunctional.

While overexpression of FtsL did not detectably suppress gp56 activity in inhibiting cell division (data not shown), overexpression of DivIC did, suggesting that additional DivIC might titrate out gp56 effects on itself and FtsL, thereby restoring their ability to effectively recruit the enzymes for septal cell wall synthesis. Recent studies from *E. coli* suggest that FtsQ/FtsL/FtsB (homologs of DivIB/FtsL/DivIC, respectively) together help bridge the activity of the earlier recruited FtsA with the later proteins needed for cell wall synthesis and invagination of the Gram-negative outer membrane (42, 43). Even without the outer membrane structure, a similar type of communication between early and late division proteins might be mediated by DivIB/FtsL/DivIC in *B. subtilis*. Consistent with this, we have isolated a spontaneous suppressor of gp56 activity that maps to *ftsA*. Though gp56 still localizes to the site of division in the presence of this FtsA suppressor, it appears that the mutant FtsA is capable of stabilizing DivIB/FtsL/DivIC components sufficiently despite gp56’s presence to still permit recruitment of Pbp2B/FtsW, similar to what is seen upon DivIC overexpression (data not shown). Alternatively, by analogy to the stronger recruitment of the key late *E. coli* divisome protein FtsN by hypermorphic alleles of FtsA (44, 45), our FtsA mutant may be able to bypass the DivIB/FtsL/DivIC requirement for recruitment of Pbp2/FtsW. The characterization of this gp56-resistant *ftsA* mutant allele and further study of the role that genes *58*-*56* might play in SP01 biology will be the focus of our future studies.

## MATERIALS & METHODS

### Bacterial Strains and Growth Conditions

All *B. subtilis* and *E. coli* strains used are listed in Table 1. *B. subtilis* strains used for experiments are derivatives of JH642 (46). *E. coli* strain AG1111 (47) was used for plasmid construction/storage and DHM1 (35) was used for bacterial two-hybrid analysis.

**TABLE 1.**
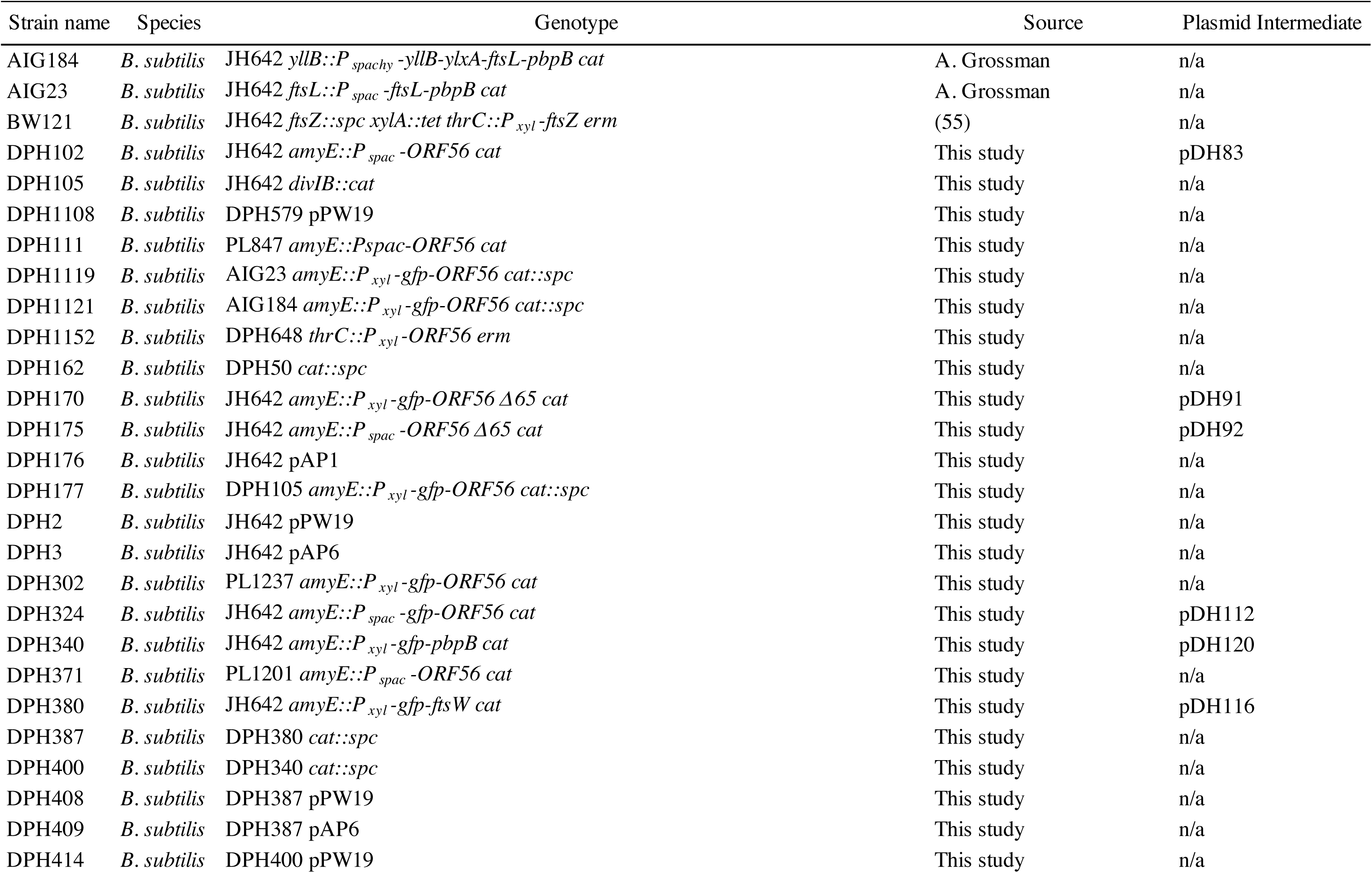
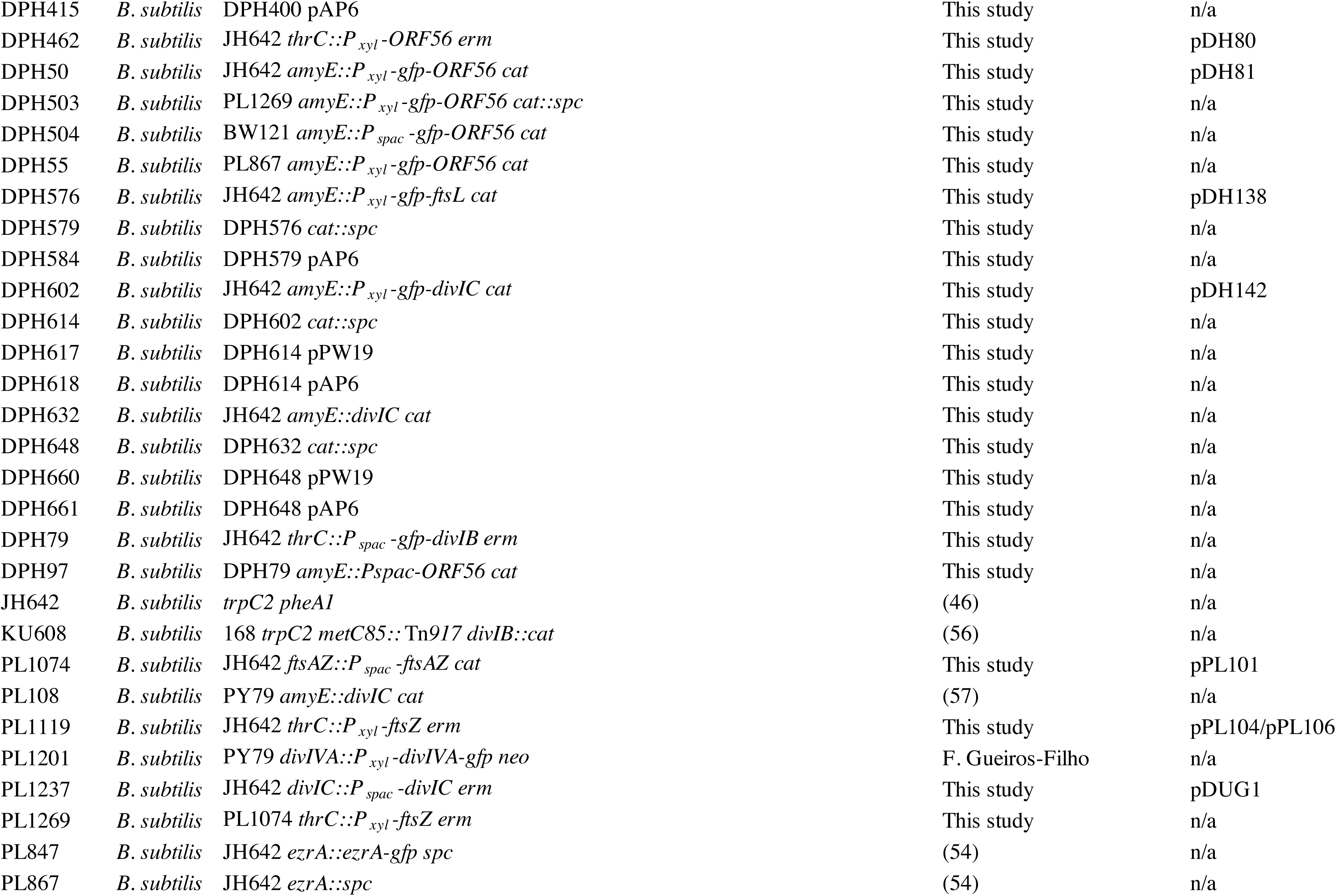
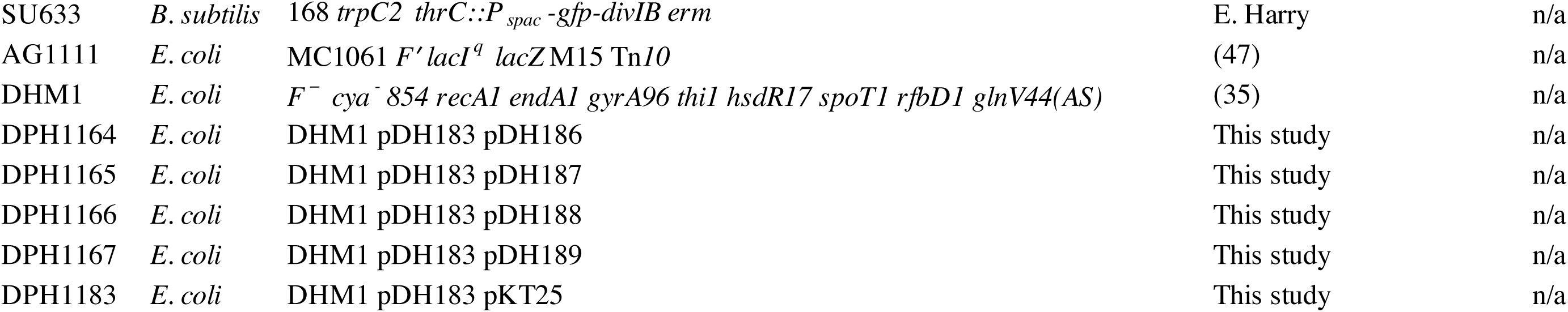
Bacterial strains used in this study.

Cells were grown in, or on, LB Lennox (1% tryptone, 0.5% yeast extract, 0.5% NaCl, 0.35 K_2_HPO_4_, pH 7.4) medium (Teknova) at 30°C for temperature sensitive strains under permissive conditions or 37°C for other strains. Sporulation was induced by exhaustion in Difco sporulation medium (DSM). Antibiotic concentrations were as previously described (23, 48). As appropriate, 1 mM Isopropyl-*β*-D-thiogalactopyranoside (IPTG) (Gold Biotechnology) was used for induction of the P_spac_ promoter or its derivatives, and 0.5% D-(+)-Xylose (Hitech) was used for induction of the P_xyl_ promoter.

For all experiments, overnight cultures of strains were diluted into fresh LB medium and were cultured to mid-exponential growth, monitored by optical density at 600 nm (OD_600_) with a Hitachi U-1800 spectrophotometer. Cultures were then diluted a second time to an OD_600_ of 0.025 – 0.05 in appropriate experimental conditions. These cultures were then grown to an OD_600_ between 0.4 and 0.6 and harvested for analysis by microscopy, fixation, or plating as described below.

### Plasmid and Strain Construction

Cloning and genetic manipulation were performed using standard techniques (49, 50) using naturally competent *B. subtilis* or chemically competent *E. coli* cells. *Pfu* DNA Polymerase (G Biosciences) was used for PCRs in an Eppendorf Mastercycler; standard restriction enzymes and T4 DNA Polymerase (New England Biolabs) were used for cloning. Plasmid DNA was prepared using the Wizard Plus SV Minipreps DNA Purification Kit, PCR and digest reactions were cleaned up using the Wizard SV Gel and PCR Clean-up System, and chromosomal DNA was prepared using the Wizard Genomic DNA Purification Kit (Promega).

The final versions of all cloning products were sequenced to verify their construction. DNA sequencing was performed by GeneWiz (South Plainfield, NJ), SeqWright (Houston, TX) or PSOMAGEN, Inc (Rockville, MD). DNA bands were visualized using a GelDoc XR Imager (BioRad) and DNA concentrations were estimated with a NanoDrop ND-1000 Spectrophotometer (Thermo Scientific).

All plasmids are listed in Table 2 and oligonucleotides purchased from IDT DNA or Fisher used for their construction are listed in Table 3. Table 2 includes details on which oligonucleotides were used for the cloning of each plasmid new to this study. Table 3 includes details on oligonucleotide sequence, indication of gene targeted for amplification, and restriction site used in the plasmid construction where appropriate. All *E. coli* strains new to this study, as well as *B. subtilis* strains DPH2, DPH3, and DPH176, were cloned by transformation with plasmids as indicated in Tables 1 and 2. All of the remaining *B. subtilis* strains new to this study were cloned by transformation with plasmids as indicated in Tables 1 and 2, followed by screening for single or double-crossover into the *B. subtilis* chromosome. Where appropriate, loss of a plasmid backbone following double-crossover was verified by antibiotic counterselection. Successful integrations at *amyE* were verified by iodine staining on starch plates, and testing for threonine auxotrophy was used to verify successful integration at *thrC*. pJL62 (51) was used for conversion of Cm^r^ Spec^s^ strains to Cm^s^ Spec^r^ ones. Strain PL1119 was created by first cloning *ftsZ* into pUC19 as indicated in Table 2 to create pPL104, followed by sub-cloning into pRDC19 to create pPL106 for transfer into the *B. subtilis* chromosome. pKT25 and pUT186 (35) were used for bacterial two-hybrid experiments (see below.)

**TABLE 2.** Plasmids used in this study.

| Plasmid name | Description | Oligos used (F/R) | Source |
| --- | --- | --- | --- |
| pAG58 | pSI-1-/pJH101-derivative with IPTG-inducible P<sub>spac</sub> for targeted integration of choice | n/a | (58) |
| pAP1 | pPW19-*ORF58-ORF57-ORF56* | n/a | (18) |
| pAP6 | pPW19-*ORF56* | n/a | (18) |
| pDH112 | pDR67-*gfp-ORF56* | oDH306/148 | This study |
| pDH116 | pEA18-*ftsW* | oDH252/253 | This study |
| pDH120 | pEA18-*pbpB* | oDH262/263 | This study |
| pDH138 | pEA18-*ftsL* | oDH292/293 | This study |
| pDH142 | pEA18-*divIC* | oDH294/295 | This study |
| pDH183 | pUT18C-*ORF56* | oDH439/440 | This study |
| pDH186 | pKT25-*ftsL* | oDH432/434 | This study |
| pDH187 | pKT25-*divIC* | oDH435/437 | This study |
| pDH188 | pKT25-*ORF56* | oDH438/440 | This study |
| pDH189 | pKT25-*ORF56* Δ65 | oDH438/440 | This study |
| pDH80 | pRDC19-*ORF56* | oDH149/148 | This study |
| pDH81 | pEA18-*ORF56* | oDH147/148 | This study |
| pDH83 | pDR67-*ORF56* | oDH149/148 | This study |
| pDH89 | pPW19-*ORF56* Δ65 | n/a | This study |
| pDH91 | pEA18-*ORF56* Δ65 | oDH147/148 | This study |
| pDH92 | pDR67-*ORF56* Δ65 | oDH149/157 | This study |
| pDR67 | pAG58-derivative integrative to *amyE* locus with IPTG-inductible P<sub>spac</sub> | n/a | (47) |
| pDUG1 | pRS14-*divIC* 5'-3' | *divIC* -5'/3' | This study |
| pEA18 | pRDC18-derivative with *spoVG* RBS followed by *gfp* and in-frame *NotI* site | n/a | (59) |
| pJL62 | pJH101-derivative for conversion of Cm<sup>r</sup> Spec<sup>s</sup> strains to Cm<sup>s</sup> Spec<sup>r</sup> | n/a | (51) |
| pKT25 | pSU40-derivative for in-frame 3'-end fusions to *cyaA* for BACTH analysis | n/a | (35) |
| pPL101 | pAG58-*ftsA* 5'-3' | *ftsA* -5'/3' | This study |
| pPL104 | pUC19-*ftsZ* | *ftsZ* -F4/R1 | This study |
| pPL106 | pRDC19-*ftsZ* | n/a | This study |
| pPW19 | pSI-1-derivative with pUB110 origin with IPTG-inducible P<sub>spac</sub> | n/a | (60) |
| pRDC18 | pRDC9-/pDG1662-derivative integrative to *amyE* locus with xylose-inducible P<sub>xyI</sub> | n/a | F. Arigoni |
| pRDC19 | pRDC9-/pDG1664-derivative integrative to *thrC* locus with xylose-inducible P<sub>xyI</sub> | n/a | F. Arigoni |
| pRS14 | pAG58-derivative with *erm* cassette in place of original *cat* cassette | n/a | A. Grossman |
| pUC19 | Standard cloning vector | n/a | (61) |
| pUT18C | pUC19-derivative for in-frame 3'-end fusions to *cyaA* for BACTH analysis | n/a | (35) |

**TABLE 3.**
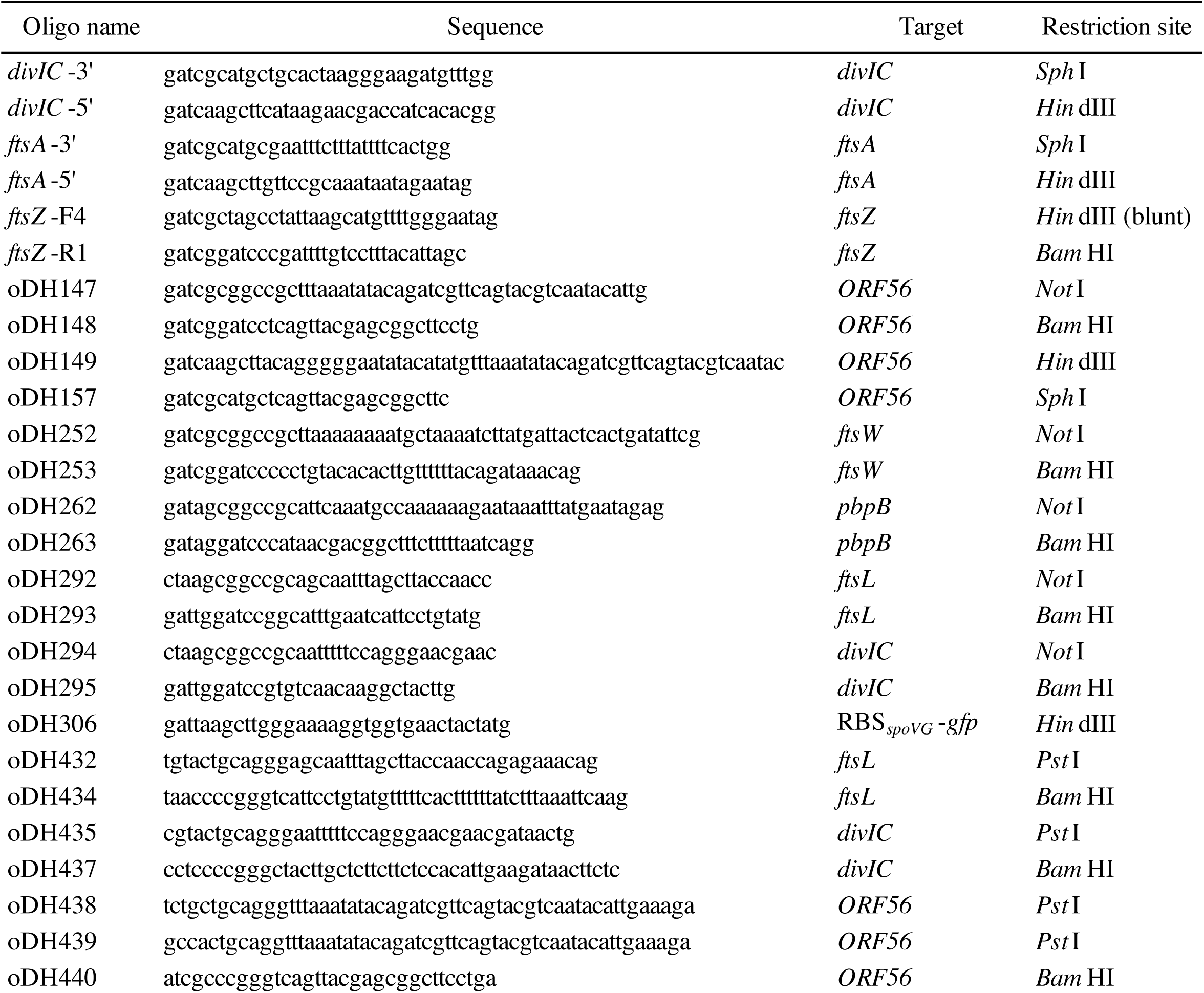
Oligonucleotides used in this study

### Spot Dilutions

As outlined above, cells used for spot dilutions were taken from fresh cultures grown to an OD_600_ between 0.4 and 0.6 under experimental conditions. Ten-fold serial dilutions of these cultures were prepared into fresh LB media (with or without IPTG as appropriate) in a 96-well plate using a multichannel pipette. A flame-sterilized and cooled, metal-pronged tool was then used to replica-plate spots of serially diluted culture onto LB plates with or without IPTG. These plates were then incubated overnight at 37°C and images were scanned using a flatbed scanner and adjusted for brightness/contrast using Adobe Photoshop.

### Cell Fixation, Microscopy, and Analysis

As outlined above, cells used for spot titers were taken from fresh cultures grown to an OD_600_ between 0.4 and 0.6 under experimental conditions and then harvested for fixation or immediate live visualization by DIC microscopy. Prior to microscopy, live cell samples were stained with the vital membrane stain FM4-64 (Molecular Probes). Cell fixation and preparation for immunofluorescence microscopy, including antibodies employed, were done as previously described (22, 23). Images were processed and analyzed for ring frequency and cell width or length (inter-septal distance for chains of *B. subtilis* cells) measurements using the ObjectJ extension (52) of ImageJ (53).

Microscopy was performed using a 100X DIC (or phase for sporulation) objective on either an Olympus BX60 microscope with a Hamamatsu C8484 camera using HC Image software (Hamamatsu), an Olympus BX51 microscope with an OrcaERG camera (Hamamatsu) using Nikon Elements Advanced Research software, or a Nikon Eclipse TE2000-E using Metavue 7.8 12.0 software (Molecular Devices). Images were processed for brightness/contrast and colorization using Adobe Photoshop.

### Bacterial Two-Hybrid (BACTH) Assays

For BACTH experiments (35), DivIC, FtsL or gp56/gp56*Δ*65 were each fused to the carboxy terminus of T25 from *Bordetella pertussis* in pKT25. Additionally, gp56 was fused to the carboxy terminus of T18 from *Bordetella pertussis* in pUT18C. Plasmids were heat-shock transformed sequentially into competent DHM1 cells and grown at 30°C. Strains were patched onto medium containing 50 g/ml X-Gal (5-bromo-4-chloro-3-indolyl-*β*-D-galactopyranoside; Gold Biotechnology), 1 mM IPTG, and antibiotics. Patches were screened for a color change following 2 days of incubation at 30°C. An image of the plate was taken using a flatbed scanner and adjusted for brightness/contrast using Adobe Photoshop.

## ACKNOWLEDGEMENTS

We thank Fabrizio Arigoni, Alan Grossman, Frederico Gueiros-Filho, Liz Harry, and Joe Lutkenhaus for the kind gift of strains and plasmids. We thank members of the Department of Microbiology and Molecular Genetics at UT Houston and the members of the Department of Biology at Canisius College for helpful discussions. This research was made possible through NIH funding to WM (GM61074) and PAL (GM127331), Frieberg Visiting Professor of Biology research funding from the Department of Biology, Washington University in St. Louis to DPH, start-up and departmental research funds from Canisius College to DPH, and Canisius Earned Excellence Program (CEEP) undergraduate student research funding to AB, JL, SA, and AM. II received partial support by the Canisius Science Scholars Program, under an NSF S-STEM award (1643649). Author contributions were as follows: Experiments: AB, II, JL, and DPH; Supporting experiments: SA, MB, DH, AM, and AP; Design, Funding, and Resources: CRS, WM, PAL, and DPH; Writing: DPH.

